# A novel R3H protein, OsDIP1, confers ABA-mediated adaptation to drought and salinity stress in rice

**DOI:** 10.1101/2021.09.20.461054

**Authors:** Liping Huang, Mohsin Tanveer, E Ji, Sergey Shabala, Mingyi Jiang

## Abstract

Abscisic acid (ABA) is a key component of many signaling networks mediating plant adaptation to various stresses. In this context, ABA-induced antioxidant defence is considered to be a main mechanism to that enhances water stress tolerance in plants. The specific details of this activation remain, however, elusive. In this work, we show that DIP1, a protein from novel R3H family, played a central role in modulating water stress tolerance in rice. *OsDIP1* transcripts were induced by hydrogen peroxide (H_2_O_2_), ABA, drought (polyethylene glycol treatment), and salt stress. Overexpression of *OsDIP1* in rice enhanced drought and salinity tolerance while knocking out *OsDIP1* by CRISPR-Cas9 editing resulted in drought and salt sensitive phenotype. The activity and gene expression of antioxidant defence enzymes, superoxide dismutase (SOD), catalase (CAT), increased in *OsDIP1*-overexpressed transgenic rice plants, while the content of malondialdehyde (MDA) decreased. In contrast, the content and gene expression of SOD and CAT, decreased, and the content of MDA increased in knockout of *OsDIP1* rice plants, suggesting that overexpression of *OsDIP1* enhances the antioxidant capacity of rice plants. The yeast two hybrid screening test revealed that OsDIP1 interacted with ZFP36, a key zinc finger transcription factor involved in ABA-induced antioxidant defence. Moreover, OsDIP1 could modulate some key ABA-responsive genes via interacting with ZFP36. Overall, our findings indicate an important role of OsDIP1 in ABA-induced antioxidant defence signaling and adaptation to salinity and drought in rice.

## Introduction

Agriculture is extremely vulnerable to climate change, with that the overall loss in the crop production from climate-driven abiotic stresses exceeding US$170Bln p.a. (Razzaq et al., 2021). Of these, drought and salinity are arguably the most critical for global food security (Liu et al., 2020). Both stresses reduced water availability to plants and cause oxidative disequilibrium that leads to oxidative damage, and in severe cases, to cell death (Baxter *et al*., 2014; Jiang and Zhang, 2001; Jiang and Zhang, 2002; Shi *et al*., 2012; Zhu *et al*., 2016a). Plants have evolved complicated mechanism to resist adversity, involving at the morphological, physiological, and molecular levels (Zhu, 2016).

Abscisic acid (ABA) accumulates quickly under water stress conditions and assist plants in their adaptation to hostile environment via plethora of signaling pathways (Jiang and Zhang, 2001; Jiang and Zhang, 2002; Shi *et al*., 2012; Baxter *et al*., 2014; Zhu *et al*., 2016a). This includes reduction in stomata aperture, activation of defence genes, and enhancement of plant antioxidant system (Raghavendra *et al*., 2010; Huang *et al*., 2018). ABA signaling pathways account for a diverse range of transcriptional, anatomical, and physiological responses. ABA-responsive dehydration responsive element binding factor, DBF, was introduced by Kizis and Pages (2002) in maize and then studied in a range of other species such as wheat (Xu et al., 2018), populus (Liu et al., 2021), and Arabidopsis (Pradeep et al., 2017). Dehydration responsive element binding (DREB) TFs comprise one of two subfamilies of the APETALA2/Ethylene Responsive Element Binding Factor (AP2/ERF) family of TFs with a single AP2 domain. Most DREB TFs bind to the dehydration-responsive element (DRE), which was initially identified in the promoter of the drought-responsive *RD29A* gene. It was demonstrated that DRE from the *RD29A* promoter is essential for gene expression induction by drought, high salinity, and low temperature (Yamaguchi-Shinozaki and Shinozaki, 1994). DREB TFs were discovered by demonstration of the fact that *Arabidopsis* nuclear protein extracts contain one or several proteins able to cause a mobility shift of oligonucleotides containing the DRE sequence (TACCGAC) in gel retardation studies (Yamaguchi-Shinozaki and Shinozaki, 1994). The first cDNAs encoding DREB proteins reported from Arabidopsis were C-REPEAT BINDING FACTOR 1 (CBF1)/DREB1 (Stockinger et al., 1997), and DREB1A and DREB2A (Liu et al., 1998). DREB TFs have subsequently been identified and characterized for a large number of plant species (Agarwal et al., 2006; Lata and Prasad, 2011; Mizoi et al., 2012; Rehman and Mahmood, 2015). Moreover, the DBF is also a part of the Apetala 2/Ethylene Response Factor (AP2/ERF) transcription factor family and induces the *rab17* (responsive to ABA) gene expression under drought stress conditions (Kizis and Pages, 2002). In maize, DBF1 and DBF2 are involved in *rab17* regulation through the drought-responsive element in an ABA-dependent pathway. Xu et al. (2008) identified three new DBF genes in *T. aestivum* (named *TaAIDFs, T. aestivum* abiotic stress-induced DBFs) by screening a wheat cDNA library after drought treatment. DREs on some promoters are involved in both ABA-dependent and ABA-independent abiotic stress signaling (Yamaguchi-Shinozaki and Shinozaki, 1994; Kizis and Pages, 2002; Agarwal et al., 2017).

A previous study found that DBF1-interactor protein 1 (DIP1) that contained two conserved core domains (R3H and SUZ) regulates the activity of DBF1 in stress responses (Saleh et al., 2006). Proteins have various conserved domains that confer their specific operating patterns. The RxxxH (R3H) domain has an ability to bind various oligonucleotides at the micromolar level without oligo (A) specificity (Bandziulis *et al*., 1989; Grishin, 1998; Liu *et al*., 2007a; Rehfeld *et al*., 2018). Removal of the R3H domain dissociated PARN (<u>p</u>oly (<u>A</u>)-specific ribo<u>n</u>uclease) into monomers, which still possessed the RNA-binding ability and catalytic functions. R3H domain is prone to dimerization, and both R3H and RNA-recognition motif (RRM) domains are essential for the high affinity of long poly(A) substrate, implying R3H domain played a critical role in the structural integrity of the dimeric PARN (Liu *et al*., 2007). While R3H domain-containing proteins are abundant in plants, their functional analysis is scarce. The R3H domain is highly conserved and widely distributed in many organisms, including eubacteria, plants, fungi, and metazoans (Saleh et al., 2006). This domain is involved in the binding of polynucleotides, including DNA, RNA, and single stranded DNA (Liepinsh et al., 2003). Moreover, the SUZ domain is a conserved RNA-binding domain found in eukaryotes and enriched in positively charged amino acids. Saleh *et al*. (2006) reported that heterologous expression of ZmDBF1 could enhance Arabidopsis drought tolerance, and DBF1-interacting protein 1 (ZmDIP1) containing an R3H domain was suggested as a potential regulator of DBF1 activity in stress responses (Saleh *et al*., 2006). In rice, the homologous protein of ZmDBF1 is OsDBF1 (OsERF48), and overexpression of *OsERF48* enhanced regulation of OsCML16 and eventually led to changes in root growth and drought tolerance (Jung *et al*., 2017). However, as OsDIP1 has 75.5% similarity with ZmDIP1 (Saleh *et al*., 2006; Liu *et al*., 2021), it remained to be answered of whether OsDIP1 is involved in rice adaptive responses to drought or some other stresses. This gap in the knowledge was filled by this study. For doing this, we have cloned OsDIP1, and investigated its subcellular localization, gene expression patterns, and impact on phenotype under water stress conditions imposed by drought and salinity. We have also surveyed OsDIP1 interacting proteins via yeast two hybrid screening method. Overall, our results indicate that OsDIP1 plays an important regulatory role as a component of the ABA-induced antioxidant defence in rice.

## Results

### 2.1 Cloning and sequence analysis of OsDIP1

The *OsDIP1* gene containing a complete open reading frame (ORF) of 1008 bp was cloned by RT-PCR. The predicted protein of OsDIP1 comprises 336 amino acids with calculated molecular mass of 37.133 kDa. *OsDIP1* encodes protein with R3H-domain and SUZ-domain (**Fig. 1A**). Structural analyses using NCBI tools have shown that OsDIP1 containing four exons and three introns ((http://rapdb.dna.affrc.go.jp/), seemed to encode two splice isoforms, and has many features of proteins dealing with RNA/DNA binding. This includes the R3H domain (Based on sequence similarities, R3H is categorized in the cysteine-rich secretory proteins, R***H, from 20 aa to 24 aa; from 72 aa to 81 aa; from 244 aa to 248 aa;) (**Fig. 1**, with black underline) and SUZ domain (from 132 aa to 178 aa) (**Fig. 1C**, with green underline), suggesting potential function at transcriptional level. OsDIP1 has two different spliceosomes implying that it can have a number of functions. OsDIP1 has high similarity with ZmDIP1, and similarity to other DIPs, showing 75.5%, 38.83%, and 35% identity respectively with ZmDIP1, AtDIP4 and AtDIP2 (**Fig. 1B**). The R3H domain is conserved among DIP proteins (**Fig. 1C**). OsDIP1, ZmDIP1, and AtDIP4 primarily distributed in group III, and was more ancient than the members of other group, suggesting that these members may have similar functions.

**Fig. 1.**
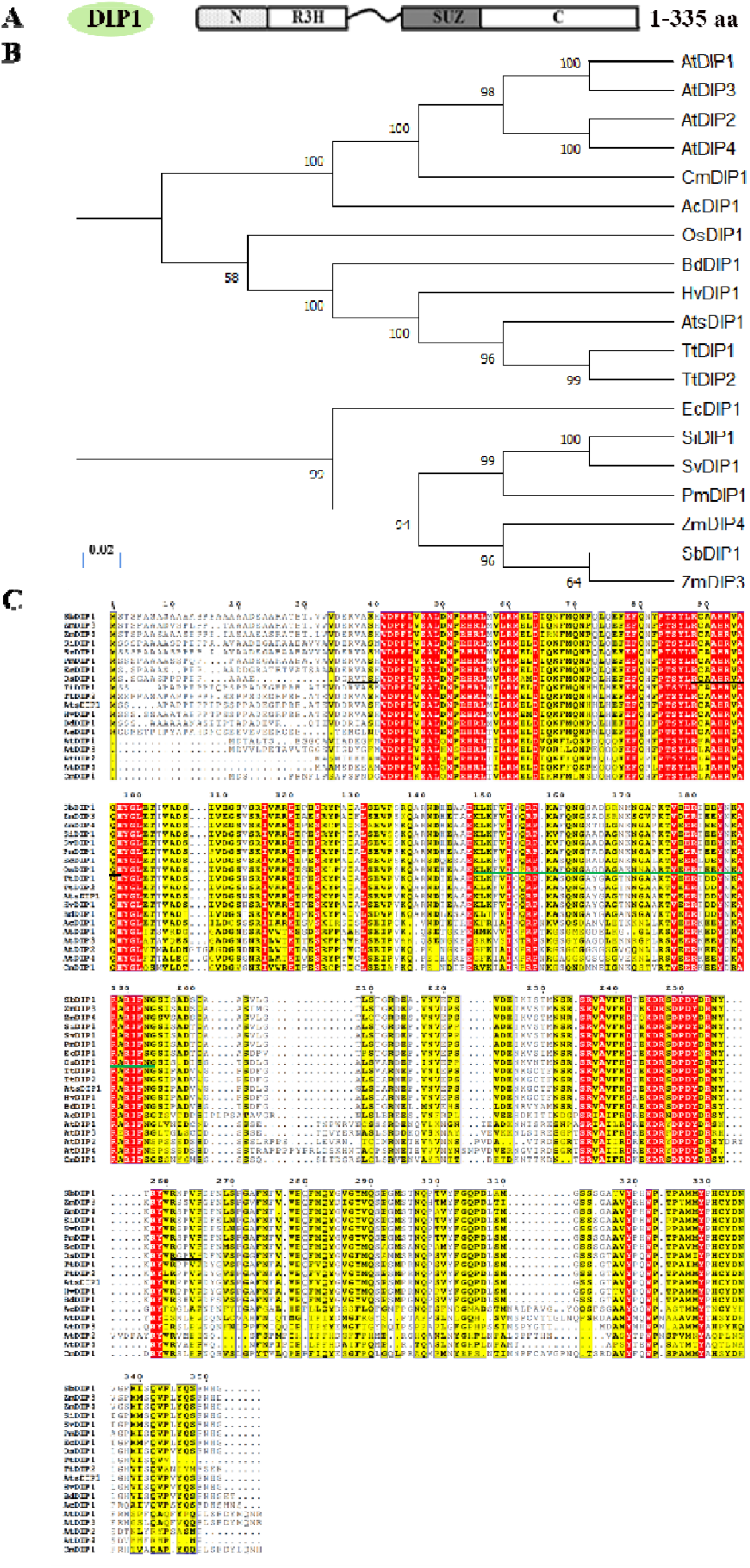
Phylogenic tree and protein sequence comparison of DBF1 interacting protein (DIPs) in key crop plants. (A) a structural diagram of OsDIP1. (B) a phylogenic tree of various DIPs in *Zea mays* (*ZmDIP1/2*), *Oryza sativa* (*OsDIP1*), *Arabidopsis thaliana* (*AtDIP1/2/3/4*), *Ananas comosus* (*AcDIP*), *Aegilops tauschii subsp. strangulata* (*AtsDIP1*), *Brachypodium distachyon* (*BdDIP1*), *Cinnamomum micranthum* (*CmDIP1*), *Eragrostis curvula* (*EcDIP1*), *Hordeum vulgare* (*HvDIP1*), *Panicum miliaceum* (*PmDIP1*), *Seteria italica* (*SiDIP1*), *Setaria viridis* (*SvDIP1*), *Sorghum bicolor* (*SbDIP1*), *Triticum turgidum* (*TtDIP1/2*) from sequence alignment of the R3H domain and SUZ domain. (C) The homologic analysis of various DIPs in (B). R***H, from 20 aa to 24 aa, from 72 aa to 81 aa, from 244 aa to 248 aa, was the R3H domain (**Fig. 1**, with black underline); The amino acids from 132 aa to 178 aa marked in green underline was the SUZ domain.

### 2.2 OsDIP1 expression pattern in tissues and under various treatments

To investigate the function of OsDIP1, tissue expression pattern via quantitative real-time PCR (qRT-PCR) and tissue chemical staining analysis were performed. qRT-PCR results revealed *OsDIP1* could accumulate in various plant tissues but was most abundant in young leaves and roots and scarcely present in old roots (**Fig. 2A**). This result suggests that *OsDIP1* may play a role in the development and function of growing leaf and root tissues.

**Fig. 2.**
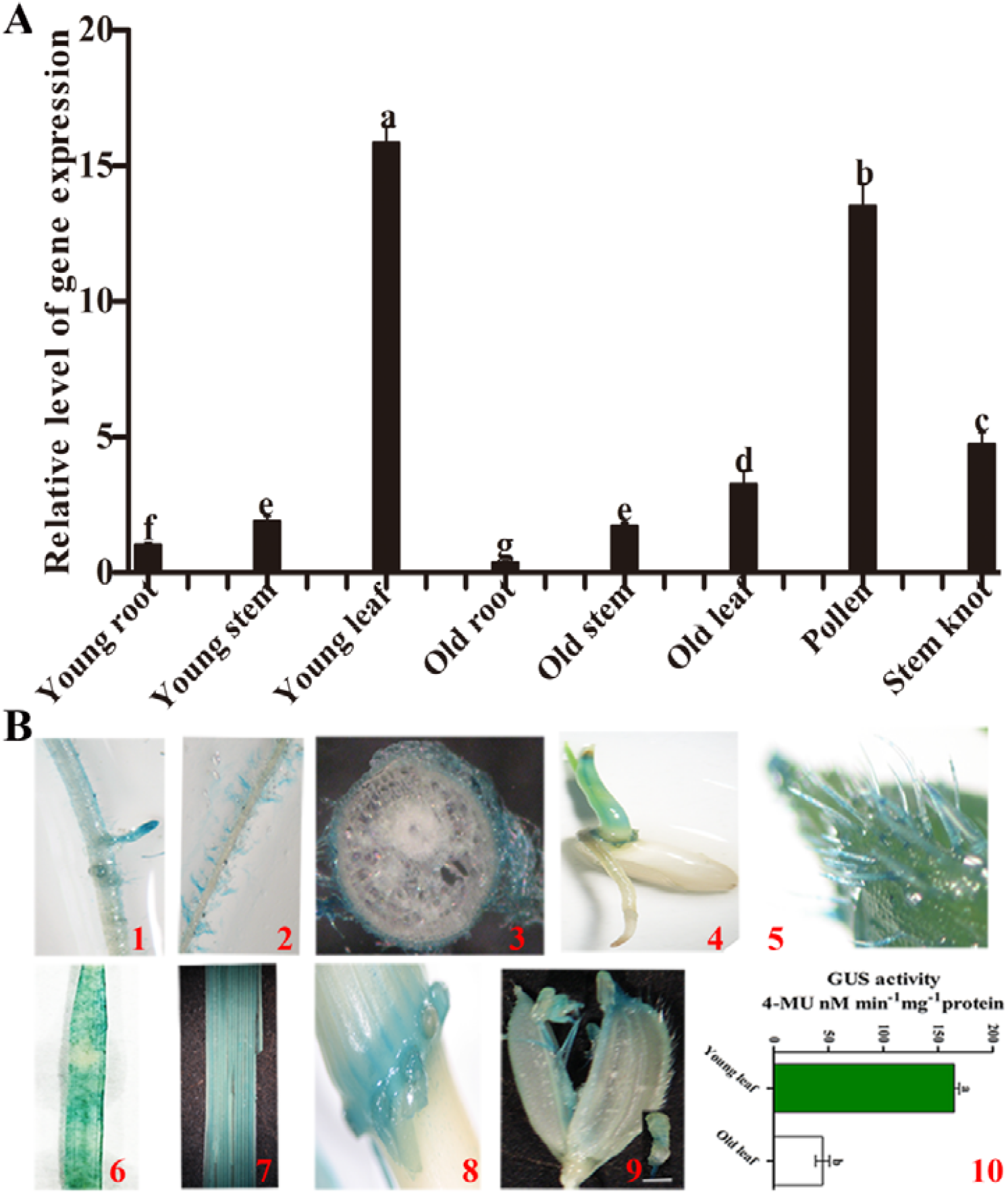
Tissue-specific expression of *OsDIP1*. (A) Tissue specific expression of *OsDIP1* by qRT-PCR (B) Histochemical GUS staining analysis. 2000 bp length of *OsDIP1* promoter was constructed to *GUS* vector and *OsDIP1-GUS* was transformed to rice. T2 generation plants were used to perform GUS staining. Representative images are shown for lateral roots (**1**); root hairs (**2**); root cross-sections (**3**); germinating seeds (**4**); arista (**5**); young and old leaves (**6, 7**); sheath and nods (**8**); and filigree and anther (**9**). Panel D**10** compares GUS activity between young leaf and old leaves. Data are mean ± SD (n = 6 to 8).

The histochemical staining assay conducted using the *GUS* reporter gene, driven by a 2,000 bp *OsDIP1* promoter region fragment, revealed that *OsDIP1* was present in most tissues including lateral roots (**Fig. 2B-1**), root hairs (**Fig. 2B-2, 3**), embryo of germinating seeds (**Fig. 2B-4**), aristas (**Fig. 2B-5**), leaves (**Fig. 2B-6, 7**), leaf sheaths (**Fig. 2B-8**), filigrees and anthers (**Fig. 2B-9**). Moreover, GUS activity analysis showed that it was obviously higher in young leaves than old leaves (a 3.5-fold; **Fig. 2B-10**), which was consistent with qRT-PCR results (**Fig. 2A**).

To link effects of ABA and H_2_O_2_ with expression of *OsDIP1* in rice plants, qRT-PCR was performed. A biphasic response was measured upon H_2_O_2_ treatment, with the first peak observed at 15 min and the second one at 180 min (**Fig. 3A**). ABA treatment caused trends qualitatively similar to that for H_2_O_2_ (**Fig. 3A**), with expression peaks also reported for 30 and 180 min. These data show *OsDIP1* displays a biphasic expression pattern after ABA treatment and H_2_O_2_ treatment.

**Fig. 3.**
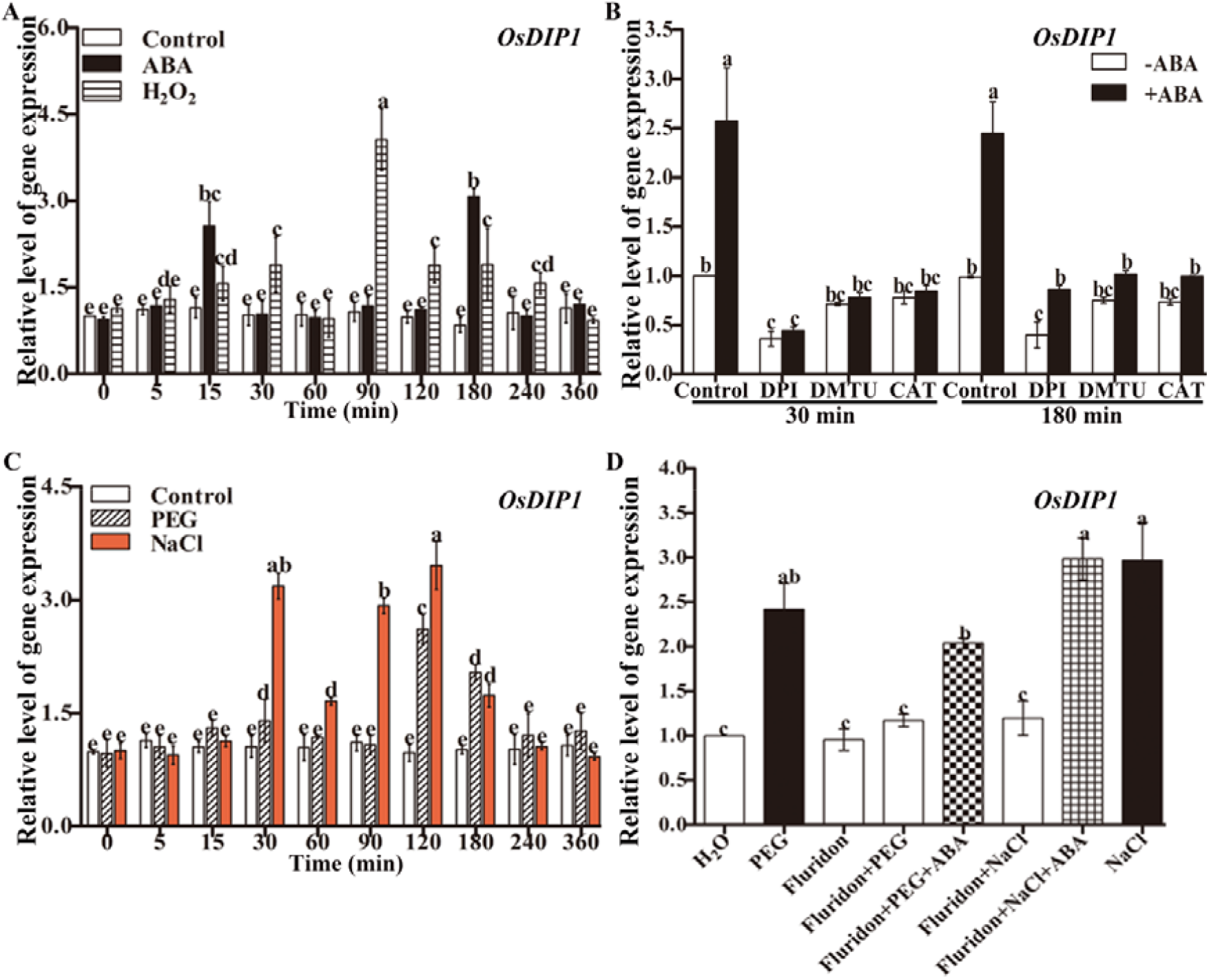
*OsDIP1* expression pattern under various stimulus. **(A)** A time course of changes in the expression of *OsDIP1* under H_2_O_2_, and ABA treatments. 10 mM H_2_O_2_, 50 μM ABA, were used to treat 10-d old rice plants. (B) The expression pattern of *OsDIP1* exposed to NADPH oxidase (a major source of the apoplastic ROS production) inhibitor diphenylene iodonium (DPI), H_2_O_2_ scavenger (dimethylthiourea, DMTU) and catalase (CAT) treated with 50 μM ABA treatment. 10 μM DPI, 100 mM DMTU, and 100 U CAT were used. (C) Time course of changes in the expression of *OsDIP1* under PEG NaCl treatments. 10% PEG6000 and 100 mM NaCl were used to treat 10-d old rice plants. (D) The expression pattern of *OsDIP1* exposed to 80 μM fluridon (an ABA inhibitor) under PEG or NaCl stress. Data are mean ± SD (n = 8).

To establish a causal link between the production of H_2_O_2_ and the expression of *OsDIP1* in ABA-signaling, rice plants were pre-treated with the NADPH oxidase inhibitor, diphenylene iodonium (DPI), which blocks the ABA-induced H_2_O_2_ production (Zhu *et al*., 2016), as well as with dimethylthiourea (DMTU), a trap for H_2_O_2_, and catalase (CAT), the enzyme eliminating H_2_O_2_. The results showed, upon modulation of H_2_O_2_ levels by these tretaments, the ABA-mediated increase in *OsDIP1* expression was significantly inhibited (**Fig. 3B)**, suggesting that H_2_O_2_ is required for ABA-induced *OsDIP1* expression.

To investigate the effects of drought and salinity stresses on the expression of *OsDIP1*, rice plants were treated with polyethylene glycol (PEG6000) and NaCl for an indicated time. As shown in **Fig. 3C**, the expression level of *OsDIP1* was significantly (2.5 to 3-fold) and rapidly induced by both drought (**Fig. 3C**) and salt treatment (**Fig. 3C**). These results indicate that OsDIP1 participates in drought and salinity stress responses. To further investigate a role of ABA in this process, fluridonet (ABA biosynthesis inhibitor) was used. Under drought or salt stress, the expression level of *OsDIP1* was significantly inhibited while treated with fluridon; this inhibition could be ameliorated by exogenous ABA treatment (**Fig. 3D**). These results indicate that OsDIP1 participation in drought and salinity stress responses is dependent on ABA.

### 2.3 OsDIP1 participates in ABA-mediated antioxidant defence

To understand the role of OsDIP1 in ABA-dependent regulation of plant redox system, two lines of *OsDIP1*-OE with an overexpressed full-length of *OsDIP1* (abbreviated OE1 and OE2) and two lines of *OsDIP1*-KO with a knock-out of *OsDIP1* by means of CRISPR-Cas9 (abbreviated KO1 and KO2) were created. As shown in **Fig. S2A**, the expression level of *OsDIP1* in OE1 and OE2 plants were significantly higher than that in WT (wild-type) plants. Two lines lacking eight base pairs (bps)- (KO1) or one (KO2) bp-deletion in the first exon of *OsDIP1* were successful knockouts (**Fig. S2B-2)**.

In the absence of ABA, there is no difference in the root length among these materials (**Fig. S3)**; however, in the presence of ABA, the root length of *OsDIP1*-OE lines was significantly shorter than that of the WT plants, whereas that of the *OsDIP1*-KO lines was longer (**Fig. S3)**. This result showed that OsDIP1 plays a role in the regulation of ABA response.

To investigate the role of OsDIP1 in plant drought and salinity tolerance, rice seedlings of OE1 and OE2, KO1 and KO2, and WT plants were treated with PEG and NaCl. Under normal growth conditions, none of above lines were significantly different from WT plants (**Fig. 4A-B**). Under drought (15% PEG 6000) and salinity (100 mM NaCl) conditions, OE1 and OE2 plants had higher survival rate, fresh weight, plant height, and root length than WT plants, while KO1 and KO2 plants performed worse than WT plants (**Fig. 4A-B**).

**Fig. 4.**
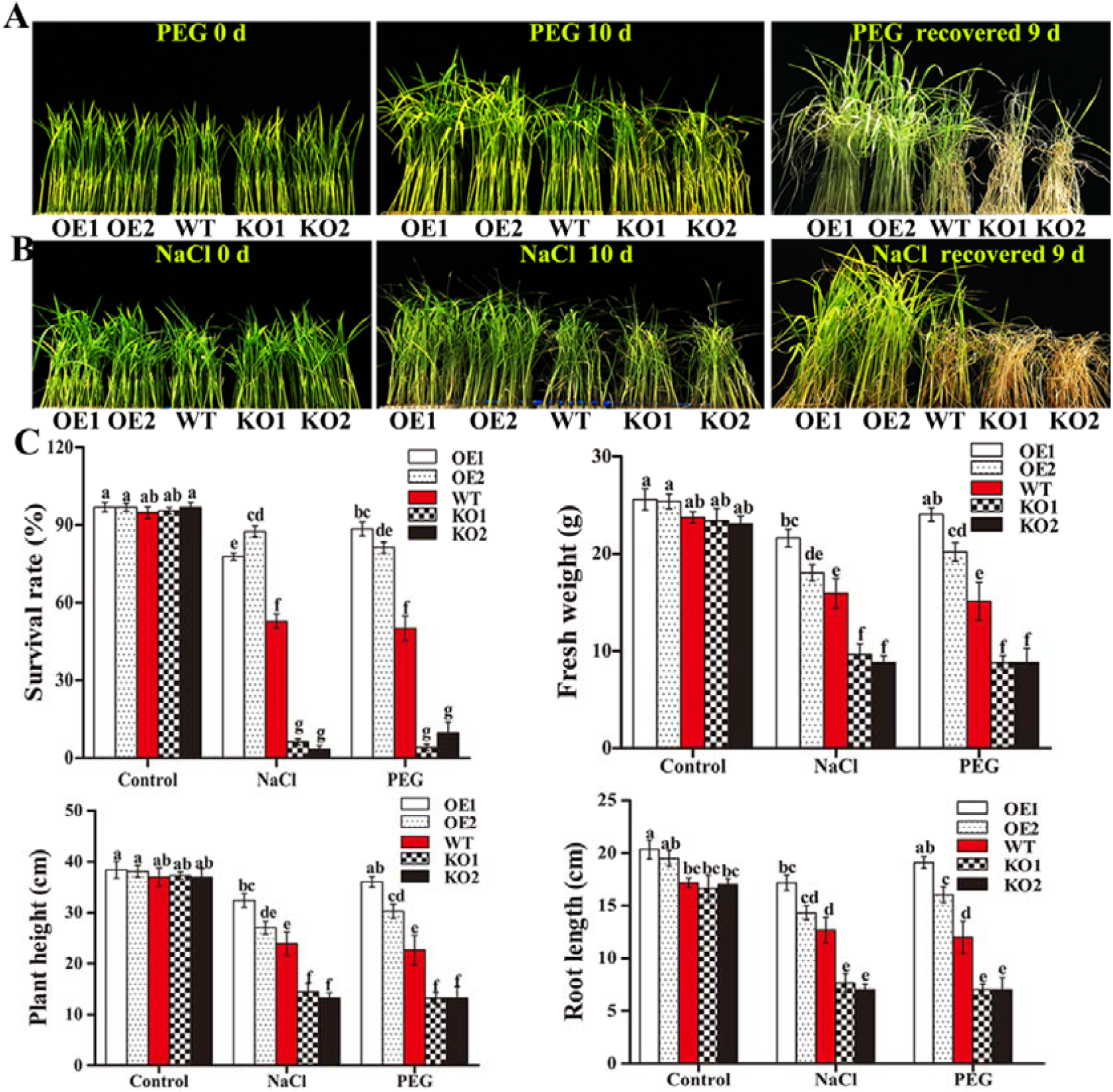
*OsDIP1* improves drought and salt stresses tolerance of rice plants. (A, B) Phenotypic analysis of responses of *OsDIP1*-OE1, *OsDIP1*-OE2, *OsDIP1*-KO1, *OsDIP1*-KO2, and WT plants grown under drought and salt stress conditions. 10-d old rice seedlings were treated with 15% PEG6000 (A) or 100 mM NaCl (B) for 10 d, and then left to recover for 9 d. Treatments were as in panels A, B. (C) The survival rate, fresh weight, plant height, and root length of various rice mutants under drought and salt stresses. 10 d-old *OsDIP1*-OE1, *OsDIP1*-OE2, *OsDIP1*-KO1, *OsDIP1*-KO2, and WT plants were treated with 15% PEG 6000 or 100 mM NaCl for 2 d. Data are mean ± SD (n = 30). Values labeled by different low-case letters are significant at *P < 0*.*05*.

To investigate the causal link between ROS production and OsDIP1 in plant adaptive responses to drought and salt stress, the differences in the expression levels of *OsRbohB/E* (a major source of apoplastic ROS production), H_2_O_2_ content, SOD and CAT enzymes activities were then investigated between OE and KO lines (Fig 5). All these characteristics were significantly higher in OE1 and OE2 than that in WT plants (**Fig. 5**) under ABA treatment; the opposite was observed for KO1 and KO2 lines. Collectively, these data implicates OsDIP1 involvement in ABA-induced antioxidant defence.

**Fig. 5.**
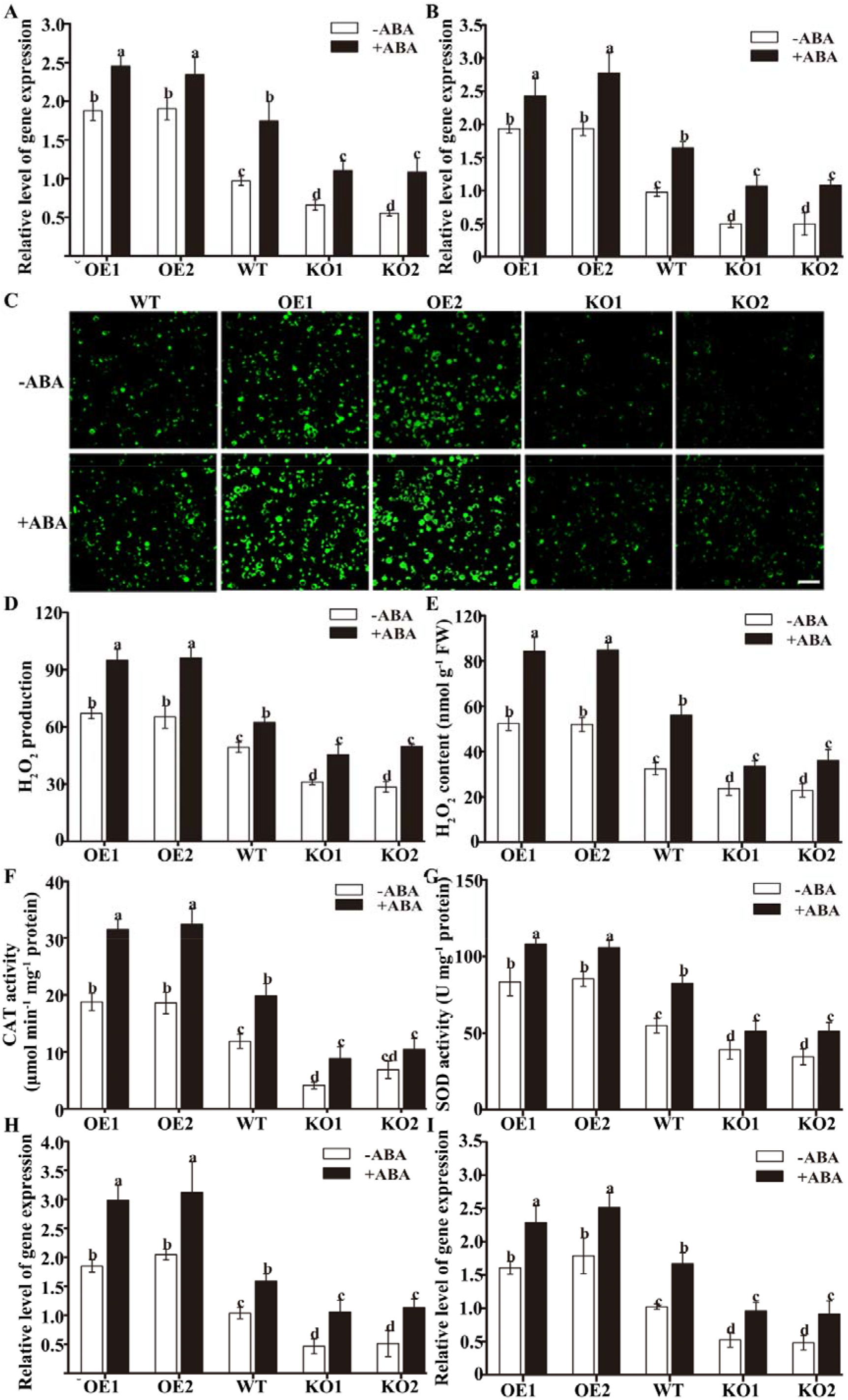
OsDIP1 participates in ABA-induced antioxidant defence. (A-B) The expression levels of OsRbohB and OsRbohE isoforms encoding NADPH oxidase in the leaves of *OsDIP1*-OE1, *OsDIP1*-OE2, *OsDIP1*-KO1, *OsDIP1*-KO2, and WT plants at various ABA levels. Rice seedlings were treated with 50 μM ABA for 30 to 90 min, and the relative expression levels of *OsRbohB/E* were determined by qRT-PCR. (C-D) Visualization of H_2_O_2_ in the protoplasts from *OsDIP1*-OE1, *OsDIP1*-OE2, *OsDIP1*-KO1, *OsDIP1*-KO2, and WT plants. Protoplasts were treated with 10 μM ABA (+ABA) or incubated without ABA for 5 min, and then loaded with H_2_DCF-DA for 10 min. H_2_O_2_ was visualized by confocal microscopy (D), and the fluorescence intensity was analysed by ZEISS LSM880 software (D). (E) - H_2_O_2_ content in the leaves of *OsDIP1*-OE1, *OsDIP1*-OE2, *OsDIP1*-KO1, *OsDIP1*-KO2, and WT plants. 50 μM ABA was given for 2 h, and the content of H_2_O_2_ in leaves was obtained. (F-I) The expression and activities of SODCc2 and CatB in the leaves of *OsDIP1*-OE1, *OsDIP1*-OE2, *OsDIP1*-KO1, *OsDIP1*-KO2, and WT plants. 50 μM ABA was given for 30 min, and the expression level of *SODCc2* (I) and *CatB* (H) were determined by qRT-PCR, the activities of SOD (G) and CAT (F) were measured. Values are means ± SD of three independent experiments. Means labelled by different low-case letters are significantly different at *P<0*.*05* according to Duncan’s multiple range test.

Further insights into this issue were obtained by comparing the ion leakage rates (**Fig. 6B**) and the content of malondialdehyde (MDA) (**Fig. 6A**) in plants upon PEG or NaCl treatments. *OsDIP1*-knock out lines displayed significantly higher ion leakage and MDA content, and lower activities of CAT and SOD under stressed conditions (**Fig. 6C, D**); in contrast, *OsDIP1*-*OE* lines had lower ion leakage and MDA content, and higher activities of CAT and SOD than WT plants (**Fig. 6C, D**).

**Fig. 6.**
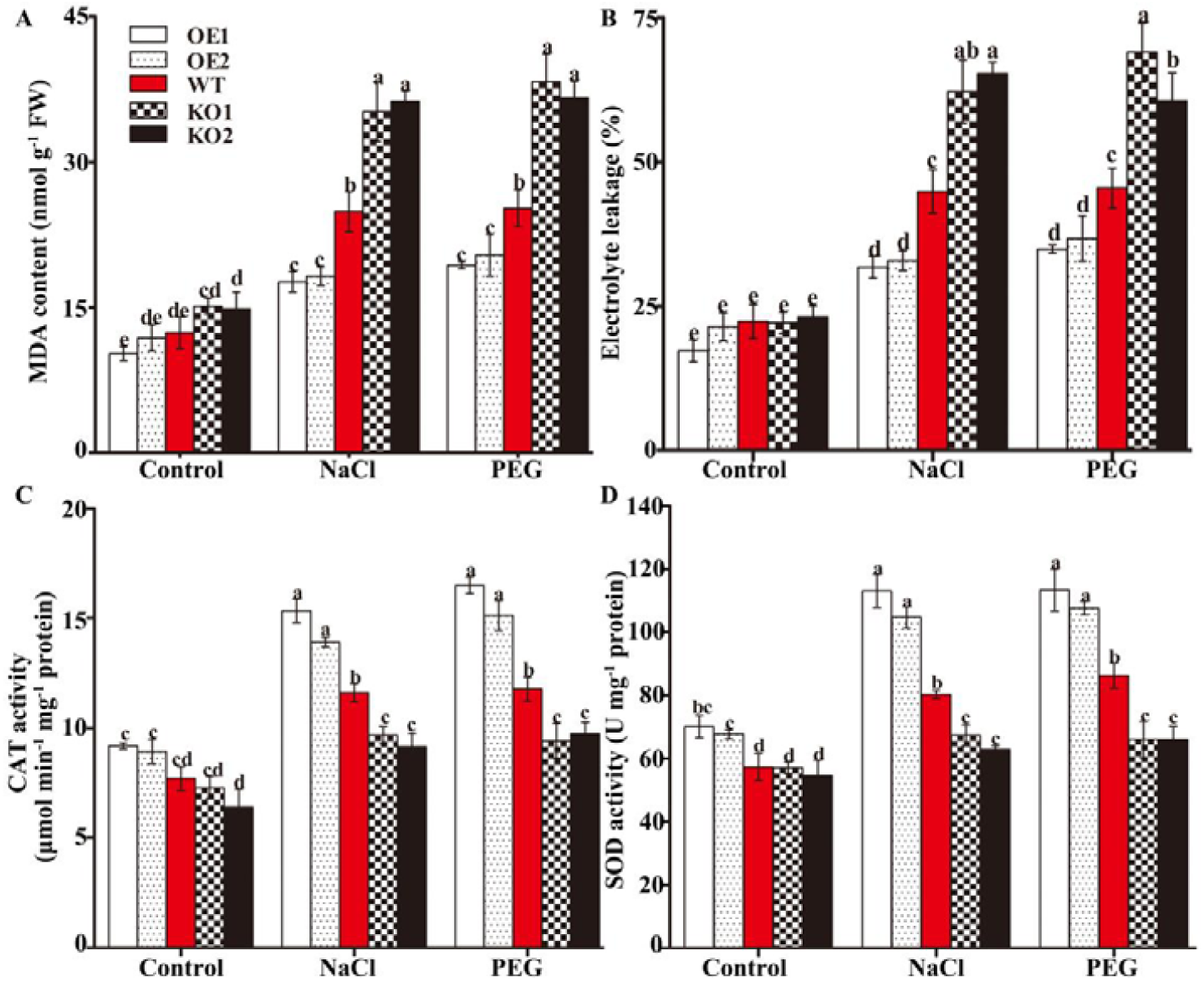
Overexpressing of *OsDIP1* enhance the antioxidant defence, while knockout of *OsDIP1* decrease the antioxidant defence under drought and salt stress. (A) MDA content and (B) electrolyte leakage (the measure of oxidative damage to lipids and plasma membranes) of various rice mutants under drought and salt stresses. 10 d-old *OsDIP1*-OE1, *OsDIP1*-OE2, *OsDIP1*-KO1, *OsDIP1*-KO2, and WT plants were treated with 15% PEG 6000 or 100 mM NaCl for 2 d. (C) catalase (CAT) and (D) SOD activity of various rice mutants under drought and salt stresses. 10 d-old *OsDIP1*-OE1, *OsDIP1*-OE2, *OsDIP1*-KO1, *OsDIP1*-KO2, and WT plants were treated with 15% PEG 6000 or 100 mM NaCl for 12 h. Data are mean ± SD (n = 30). Values labeled by different low-case letters are significant at *P < 0*.*05*.

### 2.4 Screen the interacting proteins of OsDIP1

The mechanistic basis for OsDIP1 involvement in ABA-induced antioxidant defence was further investigated in yeast two-hybrid screening experiments. One hundred and ninety-seven independent positive clones were isolated in total (**Table. S2**). Twenty-seven independent positive cDNA clones were identified as LOC_Os03g32230, designated as ZFP36 (Zhang *et al*., 2014), BsrD1 (Li et al., 2017), a zinc finger protein; seventeen were LOC_Os07g37030, cytochrome b6-f complex iron-sulfur subunit, belong to QcrA superfamily, a plastocyanin oxidoreductase iron-sulfur protein; seventeen were LOC_Os06g01210, plastocyanin, belong to cupredoxin superfamily, a type I copper protein and functions in the electron transfer from PSII to PSI; Moreover, protein argonaute 2, a central component of the RNA-induced silencing complex and related complex, etc. In total, twenty-five proteins were found to be the likely putative interacting protein of OsDIP1, implying its involvement in transcription, RNA silencing, and protein digestion.

### 2.5 ZFP36 interacts with OsDIP1 *in vitro* and *in vivo*

Previous studies have revealed that ZFP36 plays a key role in ABA-mediated antioxidant defence (Zhang et al., 2014; Huang et al., 2018). Here, yeast Y2H assay showed that yeast transformed with BD-ZFP36 and AD-OsDIP1 could grow well, while the control group didn’t grow, indicating interaction between ZFP36 and OsDIP1 **(Fig. 7A)**. *In vitro* GST pull-down assay showed that GST-ZFP36 protein bound to His-OsDIP1 did immobilize a portion of His-OsDIP1 by His antibodies, while His-OsDIP1 interaction was not observed in the control group (**Fig. 7B**). For bi-molecular fluorescence complement (BiFC)test, yellow fluorescence protein (YFP) was reconstituted when the YFP^C^-ZFP36 and YFP^N^-OsDIP1 were co-expressed in onion cells (**Fig. 7C**). In contrast, YFP fluorescence was not observed when the YFP^C^ and YFP^N^-OsDIP1 or YFP^C^-ZFP36 and YFP^N^ were co-expressed (as a control group) (**Fig. 7C**). A bright fluorescence signal was only observed in the nucleus, pointing out at it as an interaction site for OsDIP1-ZFP36 and confirming *in vivo* interaction between these two proteins. A further support comes from co-immunoprecipitation (Co-IP) assay, where His-ZFP36 was detected in a co-expression of OsDIP1-Flag and His-ZFP36 complex group with anti-His (**Fig. 7D**). In support of the above observations, subcellular localization analysis by confocal laser scanning microscopy showed that OsDIP1 was localized in the nucleus, the cytosol and the plasma membrane (**Fig. S1B**), while ZFP36 localized in the nucleus **(Fig. S1A)**. As seen in the BiFC analyses, OsDIP1 and ZFP36 co-localized in the nucleus (**Fig. S1C**) opening prospects of their direct interaction.

**Fig. 7.**
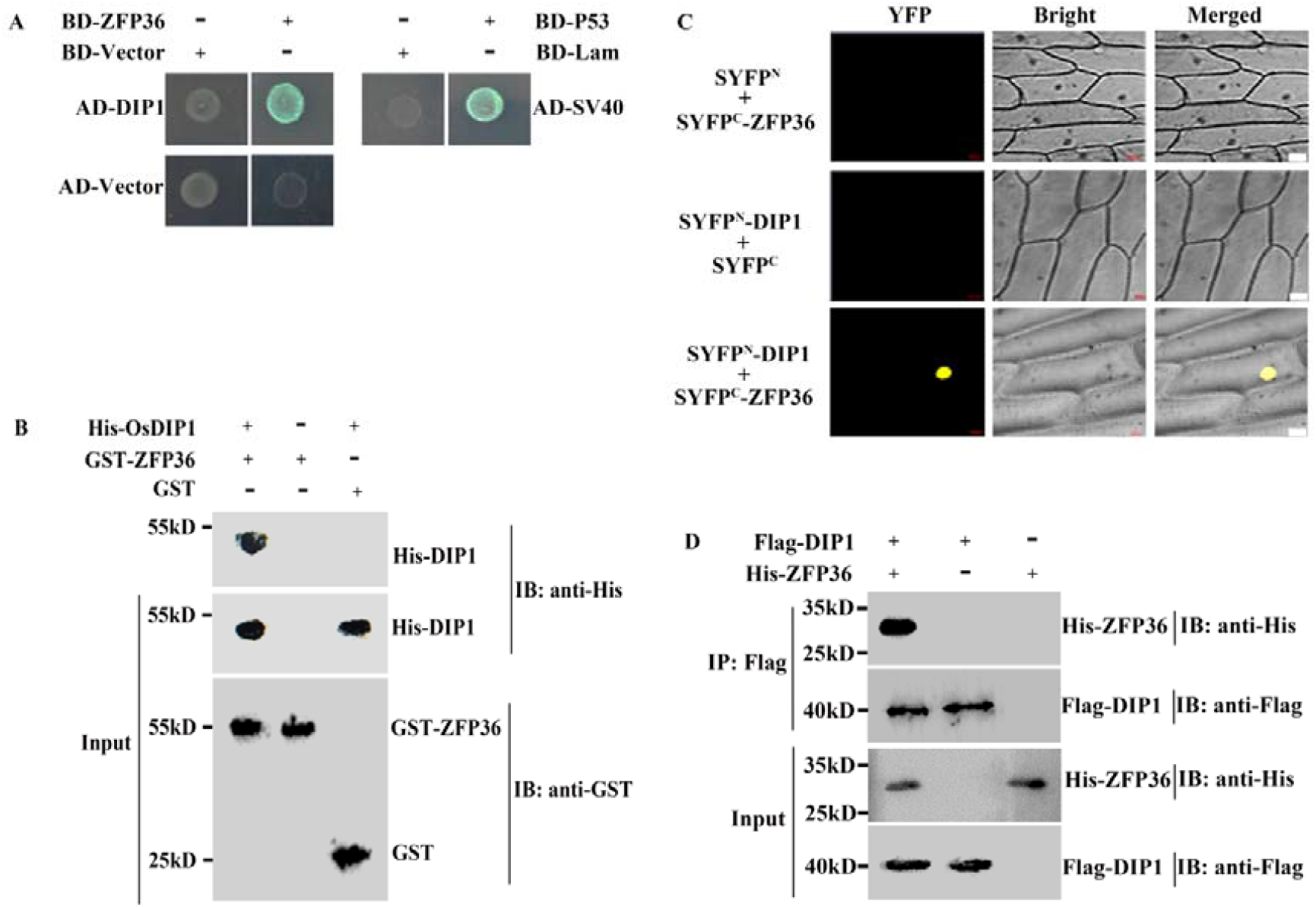
OsDIP1 directly interacts with ZFP36 *in vivo* and *in vitro*. (A) Y2H Gold yeast assay. Yeast transformants with BD-ZFP36 and AD-OsDIP1 were plated on a selective media plus X-α-gal. Unconstructed BD and AD were used as control. (B) GST pull-down test. GST-ZFP36 or GST protein were purified separately, OsDIP1-His was detected by His antibody via Western Blotting (WB). (C) BiFC assay. ZFP36 and OsDIP1 were coexpressed in onion (*Allium cepa*) epidermal cells by Gengun Bombardment. YFP signals were detected by laser confocal scanning microscope (Zeiss, LSM 880). Scale bars =50 μm. (D) Co-IP assay. His-ZFP36 and Flag-OsDIP1 were co-expressed in rice protoplast; anti-Flag was used as immunoprecipitation (IP), and the co-immunoprecipitation (Co-IP) was confirmed with anti-His.

### 2.5 OsDIP1 possibly modulate some key ABA-responsive genes to participate in ABA-induced antioxidant defence via interacting with ZFP36

Given the physical interaction of ZFP36 and OsDIP1, it might be expected that OsDIP1 interacting with ZFP36 could affect the ABA-induced antioxidant defence. To identify the role of OsDIP1 in the transcriptional activity of ZFP36, we performed LUC/REN transient expression assay. The coding sequence (CDS) region of *OsDIP1* or ZFP36 were separately recombined into *35S:YFP* vector to form effectors. Partial promoters of candidate genes - including those for ABA biosynthesis (*OsNCED1, OsNCED3*), ABA metabolism (*OsABAox2, OsABAox3*), ROS production (*RboHB*/*E*) and scavenging (*OsSODCc2, OsCatB*), and mitogen activated protein kinases (*MPK1, MPK4, MPK5, MPK7*) - were ligated to LUC/REN reporter vector (**Fig. 8A**). The corresponding effectors and reporters were transformed into rice protoplasts, and the LUC and REN enzyme activities were quantitatively detected using a Luciferase Reporter Gene Assay Kit. ZFP36 and OsDIP1 could separately enhance the promoter activities (*OsNCED1/3, OsABAox2*/*3, RboHB*/*E, OsSODCc2, OsCatB, MPK1*/4/*5/7*) than control group (**Fig. 8B-D)**; co-expressing *ZFP36* with *OsDIP1* significantly increased promoter activities as compared with separate overexpressing *ZFP36* or *OsDIP1* treatments (**Fig. 8B-D)**. ZFP36 could decrease while OsDIP1 could enhance the promoter activities of *OsABAox2*/*3* than control group (**Fig. 8C)**; co-expressing *ZFP36* with *OsDIP1* had no significant influence on the promoter activities as compared with control group (**Fig. 8C)**. Thus, OsDIP1 could enhance the transcriptional activity of ZFP36, or ZFP36 could increase the transcriptional activity of OsDIP1 to positively modulate the expression of various ABA-responsive genes.

**Fig. 8.**
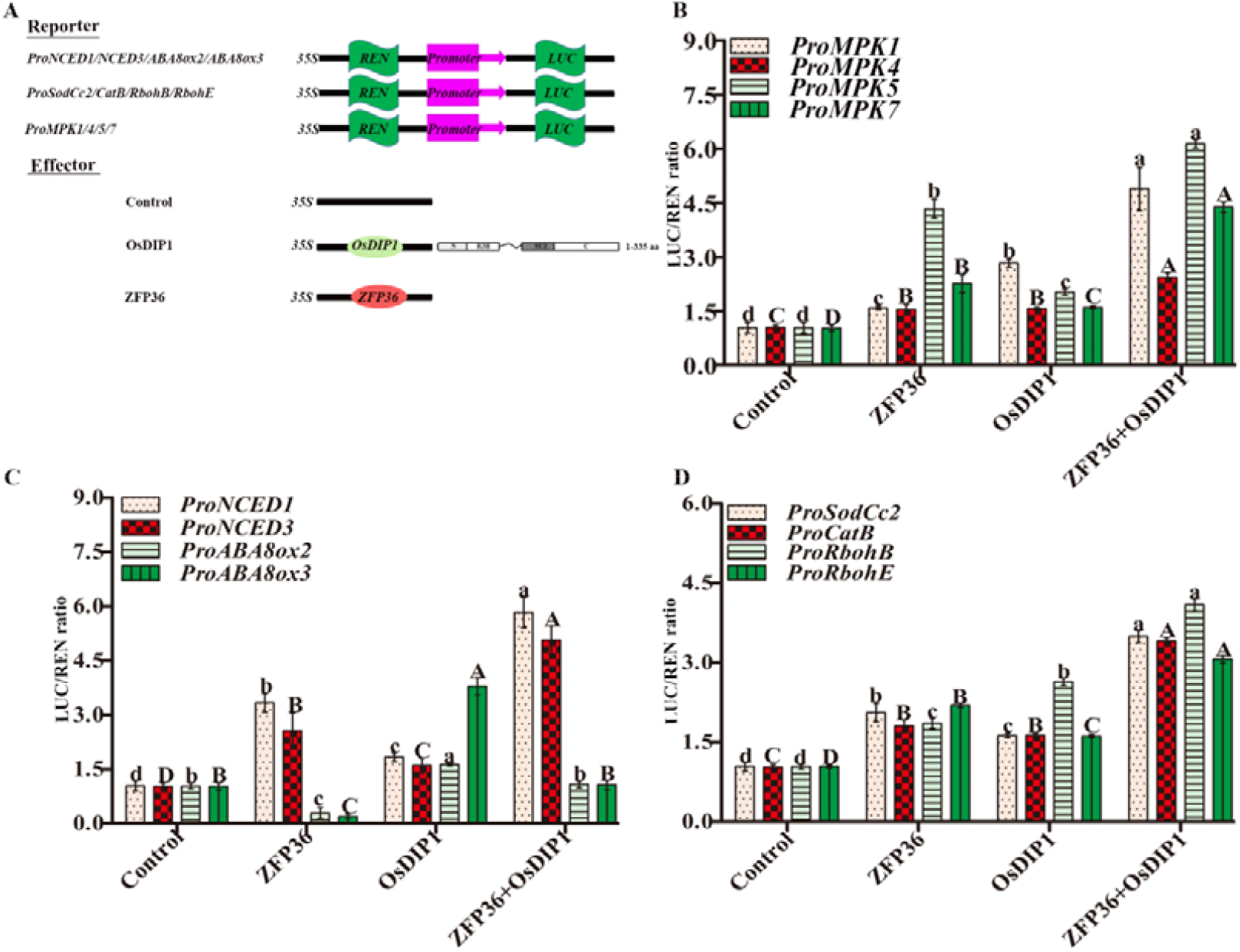
OsDIP1 alters the transcriptional activity of ZFP36. (A) A schematic diagram of the reporters and effectors for firefly luciferase (LUC)/ Renilla (REN) test in protoplasts. The promoters of *OsNCED1/3, OsABAox2*/*3, RboHB*/*E, SODCc2, OsCatB*, and *OsMPK1/4/5/7* were separately constructed to *LUC/REN* vector to form a series of reporters. ZFP36, OsDIP1, were recombined separately into *35S:YFP* vector to form various effectors. (B-D) LUC/REN assay to test the transactivation activity. Reporters and effectors in (A) were co-transformed into rice protoplasts, and the transactivation activity was determined by LUC/REN.

## Discussion

Water stress created by drought and salinity is arguably the most severe stresses affecting crop production globally. Droughts will become more frequent and severe, under current climate scenarios, and so will increase the extent of land salinization (Wani et al., 2020; Razzaq et al., 2021). Abiotic stresses adversely affect cellular homeostasis, impairing overall growth and development of plants (Choudhury *et al*., 2013; Zhu, 2016; Tanveer, 2020). Multiple physiological and molecular mechanisms act at the transcriptional and post-transcriptional level to adjust the expression of gene involved in adaptive responses to water stress. During early stages of stress, several initial stress signals activate downstream signaling processes, which subsequently activate stress-responsive mechanisms to re-establish homeostasis (Ray *et al*., 2012). Understanding the molecular mechanisms of stress tolerance in plants is, therefore, essential for sustainability and profitability of future agricultural production systems.

Abscisic acid (ABA) is known as a stress hormone that mediate a broad range of physiological and developmental responses, including those related to regulation of plant redox homeostasis and ROS signaling (Jiang *et al*., 2013; Jiang and Zhang, 2001; Jiang and Song, 2005; Hu *et al*., 2007; Hu *et al*., 2008; Huang *et al*., 2018; Shi *et al*., 2012; Shi *et al*., 2014; Zhang *et al*., 2014). Dehydration responsive element binding (DREB) factors are members of the AP2/ERF family, which comprises a large number of stress-responsive regulatory genes. The maize dehydration-responsive element (DRE)-binding factor, DBF1, is involved in the regulation of the ABA-responsive gene *rab17* through the DRE in an ABA-dependent pathway. The over-expression of the *DBF1* under the control of a stress-inducible promoter minimized the negative effect on plant growth when DBF1 was constitutively expressed (Saleh *et al*., 2006). Previous work found ZmDIP1 was mainly localized in the cytoplasm; however, after co-transformation of DBF1 and DIP1, both proteins co-localized in the nucleus. Thus, it appears that DBF1 could change the sub-localization of DIP1 for its operation (Saleh *et al*., 2006). A possibility of interaction between DIP1 and DBF1 was shown in yeast two-hybrid analyses (Saleh *et al*., 2006), but the physiological role of DIP1 has not been evaluated. Liu *et al*. (2021) found NtDIP6 can be used as a candidate gene for the molecular breeding of drought-tolerant varieties of poplar and has potential economic value in improving drought tolerance by overexpressing the gene, however, there is no more detail information of the function and mechanism of DIPs under water stress.

Here, we showed that OsDIP1 operates in ABA-induced antioxidant defence via affecting the transcriptional activity of ZFP36 thus improving rice performance under conditions of drought and salinity. The following lines of evidence support this conclusion. First, *OsDIP1* transcripts were strongly induced by drought and salinity, as well as ABA and H_2_O_2_ (**Fig. 3A, 3C**). Second, *OsDIP1*-overexpressed plants showed tolerant phenotype while *OsDIP1-*knockout rice plants showed lower tolerance to water stress than WT plants (**Fig. 4**). Third, overexpressing of *OsDIP1* in rice enhanced the ABA-induced H_2_O_2_ generation (**Fig. 5A-E**) and antioxidant defence (**Fig. 5F-I**) while knockout lines decreased it. Lastly, OsDIP1 could enhance the transcriptional activity of ZFP36 and modulate some key ABA-responsive genes to participate in ABA-induced antioxidant defence (**Fig. 8**) via interacting with ZFP36 (**Fig. 7**).

Similar to ZFP36, OsDIP1 positively regulates ABA-signaling and water stress tolerance and promotes the expression of ZFP36-modulated genes (Zhang et al., 2014; **Fig. 8**). OsDIP1 integrates plant responses to abiotic stresses by positively modulating the expression of abiotic stress-related genes. These data indicate that *OsDIP1* is a stress-responsive gene in rice, and a candidate protein for ABA-signaling via interacting with ZFP36, a key mediator in ABA-signaling.

OsDIP1 contains the R3H domain. The name of the R3H domain comes from the characteristic spacing of the most conserved arginine and histidine residues. The function of the domain is predicted to be binding ssDNA, RNA. R3H domain is prone to dimerization, and both R3H and RNA-recognition motif (RRM) domains are essential for the high affinity of long poly(A) substrate. Thus, R3H domain played a critical role in the structural integrity of the dimeric poly(A)-specific ribonuclease (PARN) (Liu *et al*., 2007). Mutational analysis of the poly (A) mRNA-binding protein (Rbs1) revealed that R3H domain was required for mRNA interactions and genetic enhancement of Pol III biogenesis (Cieśla *et al*., 2020). A R3H domain containing-like podocyte protein (R3hdml) inhibited p38 mitogen-activated protein kinase phosphorylation, and R3H domain played a key role in this process (Ishikawa et al., 2021). Further, our yeast cDNA screening test using OsDIP1 as a bait (**Table S2**) supports this notion. Many RNA processing-related proteins were isolated; one of them was Argonaute 2 (OsAGO2) protein. As this protein is the central component of the RNA-induced silencing complex, this data suggests that OsDIP1 may play a key role in the transcriptional regulation. OsAGO2 controls ROS production and increases rice susceptibility to rice black-streaked dwarf virus infection by epigenetically regulating HEXOKINASE 1 expression (Zheng et al., 2019; Wang et al., 2021). Interestingly, Argonaute 2 could interact with an RNA-binding protein MUG13.4 to mediate salinity tolerance in Arabidopsis, and MUG13.4 contains an R3H domain that mediated the above interaction (Wang et al., 2019). Taken together, it is plausible to suggest that OsDIP1 could interact with AGO2, and this process is mediated by the R3H domain. These findings highlight an important role of OsDIP1 in RNA-based silencing mechanisms to regulate the response to various stresses and prompt a call for more studies for their operation in plant adaptive responses to hostile environment.

Earlier, Vergne et al. (2008) reported that *OsDIP1* could be induced by *Magnaporthe oryzae* (rice blast fungus); OsDIP1 could also interact with Mg16820 nematode effector to resist parasitism (Naalden et al., 2018). Thus, the role of OsDIP1 appears to be broader than merely abiotic stress and could be considered as a universal regulator of plant adaptive responses to both biotic and abiotic stresses. Moreover, a mitogen activated protein kinase kinase, OsMKK6 (LOC_Os01g32660) could interact with OsDIP1 (Ding et al., 2009; Rohila et al., 2006). OsMKK6 is highly inhibited exposed to rice black-streaked dwarf virus infection and plays a key role in plant-pathogen interaction (Mohamed et al., 2017). Meanwhile, it is also regulated by salt and drought stresses (Kumar et al., 2008). Thus, OsMKK6 plays a key role in biotic and abiotic stresses, implying the likely role of OsDIP1 in modulating biotic and abiotic stress via interacting with OsMKK6 or other proteins. Here, we also find OsDIP1 interacts with ZFP36 *in vitro* and *in vivo*. ZFP36 belongs to C_2_H_2_-type zinc finger proteins. These proteins are abundant in plants, with almost 0.7% of C_2_H_2_-type ZFPs reported in *Arabidopsis* genome, about 0.8% in yeasts, and 0.7% in rice. Most of the C_2_H_2_-type ZFPs in plants have a highly conserved QALGGH sequence in zinc finger domains, and this kind of ZFPs are referred to as Q-type ZFPs (Alam *et al*., 2019). Among these subfamilies, C_2_H_2_-type ZFPs were widely explored in plant growth, development, and signal transduction under stresses, including in model plants such as *Arabidopsis thaliana, Triticum aestivum, Glycine max*, and *Oryza sativa*. Various C_2_H_2_ zinc finger proteins can regulate ABA accumulation in plant tissues, thus conferring plant responses to abiotic stress (Luo et al., 2012; Wang et al., 2016; Figueiredo et al., 2012; Wang et al., 2019; Han et al., 2020). In rice, ZFP36 has been shown to play a pivotal role in ABA-induced antioxidant defence (Figueiredo et al., 2012; Zhang et al., 2014; Huang *et al*., 2018a;b), and it has also been reported to negatively regulate rice disease tolerance (Li et al., 2017). Thus, it appears that OsDIP1 plays an essential role in modulating plant responses to biotic and abiotic stress, and execute this via interacting with ZFP36.

In conclusion, our result revealed OsDIP1 could enhance the transcriptional activity of ZFP36, or ZFP36 could increase the activity of OsDIP1 to positively modulate the expression of various ABA-responsive genes to participate in ABA-mediated antioxidant defence.

## MATERIALS AND METHODS

### Plant materials and growing conditions

Rice (*Oryza sativa* L., Nipponbare cultivar) plants were used as the wild type (WT) plants or to generate various mutant plants. Rice seeds were imbibed on moist filter paper at 28°C in the dark for 72 h, and rice seedlings were grown under the conditions as reported previously (Huang et al., 2018). When the second leaves were fully expanded, the plants were collected and used for further investigations.

### RNA isolation and qRT-PCR

Rice leaves were ground into powder by adding liquid nitrogen. The total RNA was extracted with RNAiso produced by TaKaRa (TaKaRa, Dalian, China), according to the instruction manual. Quantitative real-time PCR (qRT-RCR) assay was completed in CFX96 made by Bio-Rad. *OsActin1* (Os03g0718100) was used as an internal reference gene. Quantitative results were analyzed using the 2^-^□□^Ct^ method. Experiments were repeated at least three times.

### Subcellular localization

For subcellular localization of OsDIP1 and ZFP36, the two genes of coding sequence were separately fused to yellow fluorescent protein (YFP) at the N-terminus driven by *35S* promoter (*35S*:OsDIP1-YFP, *35S*:ZFP36-YFP) and native promoter (*OsDIP1*:OsDIP1-YFP, *ZFP36*:ZFP36-YFP) with *Bam*HI and *Kpn*I. These plasmids were transiently expressed in rice protoplast, and *35S*: YFP plasmid was used as control. The ZEISS LSM 800 (ZEISS, Germany) microscope was used to detect fluorescence.

For co-localization test, we connected ZFP36 to red fluorescent protein (mCherry) with *Bam*HI and *Kpn*I to construct ZFP36-mCherry. Then ZFP36-mCherry and OsDIP1-YFP were transiently co-transformed into onion cells by GenGun bombardment. The ZEISS LSM 800 (ZEISS, Germany) microscope was used to detect fluorescence.

### Physiological Measurements

We generated transgenic rice plants, in the background of Nipponbare cultivar. Overexpression of the full-length of *OsDIP1* (*OsDIP1*-OE), knock-out of the *OsDIP1* mediated by CRISPR-Cas9 (*OsDIP1*-KO) were generated by Biorun Biotechnology Company (Wuhan, China).

To obtain *OsDIP1*-OE, the full-length open reading frame (ORF) was fused to the plant binary vector pSUPER1300 under the control of CaMV *35S* promoter with *Xba*I and *Kpn*I to generate pSUPER1300-*OsDIP1*. The recombined plasmids were transformed into rice via *Agrobacterium*-mediated transformation method. Homozygous T3 seeds were harvested and used for further test. For *OsDIP1*-KO plants, CRISPR/Cas9 system from Biorun Biotechnology Company (Wuhan, China) was used. The sgRNA (GTAGCAGATAGCTTAGTAGA<u>TGG</u> (**Table S1**) was produced in the pRGE6 vector containing Cas9 via *Agrobacterium*-mediated transformation method. To test the CRISPR/Cas9 *in vivo*, genomic DNA was isolated by hexadecyltrimethylammonium bromide (CTAB) solution, the primers (**Table S1**) flanking the designed target site was used to determine it. Two independent lines (OE1 and OE2) with *OsDIP1-*overexpression, and two independent lines (KO1 and KO2) with *OsDIP1*-knock out were selected for following analysis.

To evaluate the phenotype of the transgenic and mutant rice plants under water stresses, 10-d old rice plants were treated with either 15% PEG 6000 (mimicking drought) and 100 mM NaCl (salinity) for the indicated time, and the survival rates of the rice plants were counted after recovery at normal conditions for 9 d. For analyzing the oxidative damage to lipids and plasma membranes, plants were treated with 15% PEG 6000 and 100 mM NaCl for 2 d, and the content of malondialdehyde (MDA) in leaf segment homogenates, prepared in 10% trichloroacetic acid containing 0.65% 2-thiobarbituric acid (TBA), and heated at 95 °C for 20-30 min, was determined according to the instructions of MDA test kit (purchased from Beyotime Biotechnology, Beijing, China). Electrolyte leakage was assayed using electrolyte leakage test kit (Beyotime Biotechnology, Beijing, China) as described by Jiang and Zhang (2001). For root growth test, 3-d-old seedlings were individually transferred to 1/2 MS medium with different concentration of ABA (0, 5, and 10 μM), then were further incubated in the growth chamber for another 15 d, the length of primary roots was measured. H_2_O_2_ content in protoplasts was determined by H_2_O_2_ probe 2′,7′-dichlorofluorescein diacetate (H_2_DCF-DA). Rice protoplasts were obtained by isolating cells from rice mutants. The H_2_O_2_ probe was loaded by incubating protoplasts for 15 min. Briefly, 50 μL of supernatants and 100 μL of test solutions were added and mixed in the test tubes at 30 °C for 30 min, and the optical density of solution was measured immediately with a LSM800 spectrometer at a wavelength of 560 nm. The content of H_2_O_2_ was quantified from absorbance readings using the calibration curve. H_2_O_2_ content in transgenic plants was determined using the Hydrogen Peroxide Assay Kit test kit (Beyotime Institute of Biotechnology, Shanghai, China). For the analysis of antioxidant enzymes, rice plants were treated with 50 μM ABA for indicated time, and the genes expression and activities of superoxide dismutase (SOD) and catalase (CAT) were measured using a standard method (Jiang and Zhang, 2001).

### GUS staining

The histochemical analysis followed the method described by Ai *et al*. (2009). Briefly, we generated transgenic plants expressing β-glucuronidase (GUS) driven by the native promoter of *OsDIP1* (*pOsDIP1-GUS*). The promoter region of *OsDIP1* was constructed to *pCAMBIA1301::GUS* vector to form the recombinant plasmid *OsDIP1*::GUS. The plasmid then was transformed into rice callus to obtain *OsDIP1*::*GUS* transgenic plants. T2 transgenic seeds were used for GUS staining by incubation at 37 °C for 4-6 h in 0.5 M phosphate buffer (pH 7.2), which contained 100 mM K_4_Fe(CN)_6_, 100 mM K_3_Fe(CN)_6_, 10% v/v Triton X-100, 0.5 M EDTA, and 0.5% w/v X-Gluc. After dyeing, the tissue was placed in an incubator at 37°C and decolorized in anhydrous ethanol, and pictures were taken with a Canon camera.

### Y2H Screening

The full-length of OsDIP1 was constructed into the *pGBKT7* vector (Clontech, TaKaRa, Dalian, China) as the bait. Total RNA was isolated from rice leaves treated with 50 μM ABA, and a rice cDNA library was produced as described previously (Ni et al., 2019). The bait plasmid was transformed into Y2H Gold yeast strain, and the cDNA library was transformed into Y187 yeast strain, following the method described in Clontech’s yeast protocols handbook. Screening of interaction clones was performed as described by the manufacturer’s instructions (Clontech, TaKaRa, Dalian, China). The screened transformants from the cDNA library were transferred to grow on the stringent dropout medium which lack of Tryptophan (Trp), Leucine (Leu), Histidine (His), and Adenine (Ade) (SD/-Trp-Leu-His-Ade). The library plasmid responsible for the activation of reporters was amplified by PCR and was cloned into *pGADT7*, then were transformed into Y187 yeast strain, the one-on-one interaction was identified by yeast two-hybrid assay.

### Yeast two-hybrid assay

Yeast strains Y2H and Y187 were activated on YPDA medium, the yeast two-hybrid assay was carried out following the manufacture as described in Clontech’s yeast protocols handbook. Briefly, an appropriate amount of a single colony was transferred to 5 mL liquid YPDA medium and allowed to growth until OD_600_ reached 2.0. The strain was then activated again with a liquid YPDA medium, until OD_600_ reached to 0.4-0.6, and collected material was used for subsequent experiments. Before the experiment, ZFP36 was connected to *pGBKT7* vector to form BD-ZFP36 and OsDIP1 to *pGADT7* to form AD-OsDIP1. BD-P53, BD-Lam and AD-SV40 were used as the control group, where BD-P53+AD-SV40 combination was the positive control, and BD-Lam+AD-SV40 was the negative control. Using the principle of yeast heterologous fusion, two transformed strains were mixed and cultured, and then selected on SD/-Trp-Leu, for preliminary screening. The screened colonies were transferred to the solid quadra-deficient culture medium, SD/-Trp-Leu-His-Ade. Additionally, a moderate concentration of X-α-Gal was used to further test the interaction.

### Gst Pull-down assay

*Escherichia coli* (*E. coli*), strain BL21, purchased from CWBIO company (Suzhou, China), was used to express proteins. The CDS region of *ZFP36* and *OsDIP1* were cloned into *pGEX-4T-1* (a carrier labeled with GST protein) with *Bam*HI and *Eco*RI, and pET-30a (a vector tagged with His protein) with respectively, to form GST-ZFP36 and OsDIP1-His structures. The fusion plasmids were imported into *E. coli* BL21. The transformed BL21 was cultured in a liquid LB medium containing corresponding antibiotics to an appropriate concentration, then it was transferred to 200 mL medium and cultured for additional 4 h. When the concentration of *E. coli* reached OD_600_=0.60, 0.50 mM isopropyl β-D-thiogalactoside (IPTG) was added to the medium. Bacteria were collected and lysed with an ultrasonic crusher. The supernatant was purified with Promega™ His- and GST-magnetic beads. OsDIP1-His was separated from the magnetic bead with 500 mM imidazole, and GST-ZFP36 was purified and bound to GST-magnetic beads. 100 mg of each protein were mixed together in a GST pull-down buffer (150 mM NaCl, 1 mM EDTA, 1 mM DTT, 0.5% NP-40, 50 mM Tris-HCl, pH=7.4), incubated at 4□ for 2 to 4 h with shaker. The magnetic beads were then washed 5 times with PBS buffer. Western blot (WB) test was used to verify the interaction.

### Co-IP assay

The coding sequence of *OsDIP1* was inserted to *35S:YFP* vector with Flag-tag to obtain OsDIP1-Flag, while *ZFP36* was constructed to *35S:YFP* vector with His-tag to form His-ZFP36. His-ZFP36 and OsDIP1-Flag were transiently co-expressed in rice protoplast, and two plasmids were separately transformed into rice protoplast as the control group. Two-week old rice seedlings were used to produce protoplasts using the procedure described by Zhang et al. (2011) with minor modification. Briefly, K3 buffer containing cellulase, macroenzyme, and D-Mannitol, was used to configure the enzymatic hydrolysate. Plant tissue with chopped with the double-side blade and soaked in the K3 buffer and vacuumed for 1 hour in dark. The mix was centrifugated at 100-300 rpm for 4 hours. Tissue debris were cleaned and filtered with W5 buffer (154 mM NaCl, 125 mM CaCl_2_, 5 mM KCl, 2 mM MES, pH 5.7), to isolate pure protoplasts. Recombinant plasmids were transformed with PEG buffer system (40% PEG 4000, 0.4 M D-Mannitol, 100 mM CaCl_2_). Transformed protoplasts were cultured for 16 h, then were harvested to extract total protein. Flag antibody was used to immunoprecipitate the OsDIP1-Flag and His-ZFP36 complex, His antibody was used to WB detection.

### BiFC

BiFC was performed in onion epidermal cells. The recombinant plasmids *SYFP*^*C*^*-ZFP36* and *SYFP*^*N*^*-OsDIP1* were obtained by recombination of *SYFP*^*C*^ and *SYFP*^*N*^ empty carriers with the CDS of *ZFP36* and *OsDIP1*, respectively. The inner epidermis of onion was separated with forceps on MS medium. Plasmids were diluted to 1.0 mg/μL and assayed as per Lai *et al*. (2013) with minor modification. 8.5 μL gold powder and 1 μL plasmid were collected into a 1.5 mL Eppendorf tube and mixed with a Vortex for 60 s. 7.5 μL of 2.5 M CaCl_2_ and 3 μL of 0.1 M spermine were added and mixed. The mixture was centrifuged with 12000 rpm speed and the supernatant was discarded. The precipitate was washed twice with 80 μL anhydrous ethanol, and finally suspended with 10 μL anhydrous ethanol. The completed sample was dropped to the center of the carrier membrane and injected into onion cells using Biolistic Technology. Fluorescence was detected by the laser confocal technique (ZEISS LSM 800, ZEISS, Germany), with the excitation light being 488 nm.

## Abbreviations

ABA: Abscisic acid
APX: ascorbate peroxidase
CAT: catalase
DAPI: 4’,6-diamidino-2-phenylindole, dihydrochloride
DMTU: dimethylthiourea
DPI: diphenylene iodonium
*E. coli*: *Escherichia coli*
H_2_DCF-DA: 2′,7′-dichlorofluorescein diacetate
H_2_O_2_: hydrogen peroxidase
NCED: (9-cis-epoxycarotenoid dioxygenase)
LUC/REN: firefly luciferase/renilla
MDA: malondialdehyde
NaCl: sodium chloride
PEG: polyethylene glycol
Rbohs: respiratory burst oxidase homologs
SOD: superoxide dismutase
YFP: yellow fluorescence protein
ZFP: zinc finger protein.

## Acknowledgements

We are grateful to Prof Wenhua Zhang from Nanjing Agricultural University for supplying pSUPER1300 and pCAMBIA1301 vectors, and Prof. Hongxuan Lin from Shanghai Institute of Plant Physiology and Ecology, Shanghai Institutes for Biological Sciences, Chinese Academy of Sciences for providing rice protoplast transformation vectors to study the transactivation of TFs.

## Funding

This work was supported by the National Natural Science Foundation of China (grants 31901202, 31672228), National Distinguished Expert Project (WQ20174400441), Project for High-level Talents of Foshan University (gg07102, gg05003/071), and by the Australian Research Council (DP150101663).

## Conflict of interest

The authors have declared no conflict of interest.

## Accession Numbers

Sequence data from this article can be found in the GenBank/EMBL data libraries under the following accession numbers: OsDIP1, LOC_Os05g34070, ZmDIP1/2, GRMZM2G010302/GRMZM2G055970, AtDIP1/2/3/4, AT2G40960/AT3G10770/AT3G56680/AT5G05100.1; ZFP36, LOC_Os03g32230; APX1, LOC_Os03g17690; OsSODCc2, LOC_Os03g22810; GAPDH, LOC_Os02g38920; Actin1, LOC_Os03g50885. AcDIP1, OAY67162.1; *AtsDIP1*, XP_020201064.1; *BdDIP1*, XP_003568455.1; *CmDIP1*, RWR79788.1; *EcDIP1*, TVU19363.1; *HvDIP1*, KAE8789012.1; *PmDIP1*, RLN00811.1; *SiDIP1*, XP_004962092.1; *SvDIP1*, XP_034586427.1; *SbDIP1*, KAG0518079.1; *TtDIP1/2*, VAH07401.1/VAH19183.1; *ZmDIP3/4*, NP_001105871.2/NP_001148805.2.

---

**Fig. S1.**
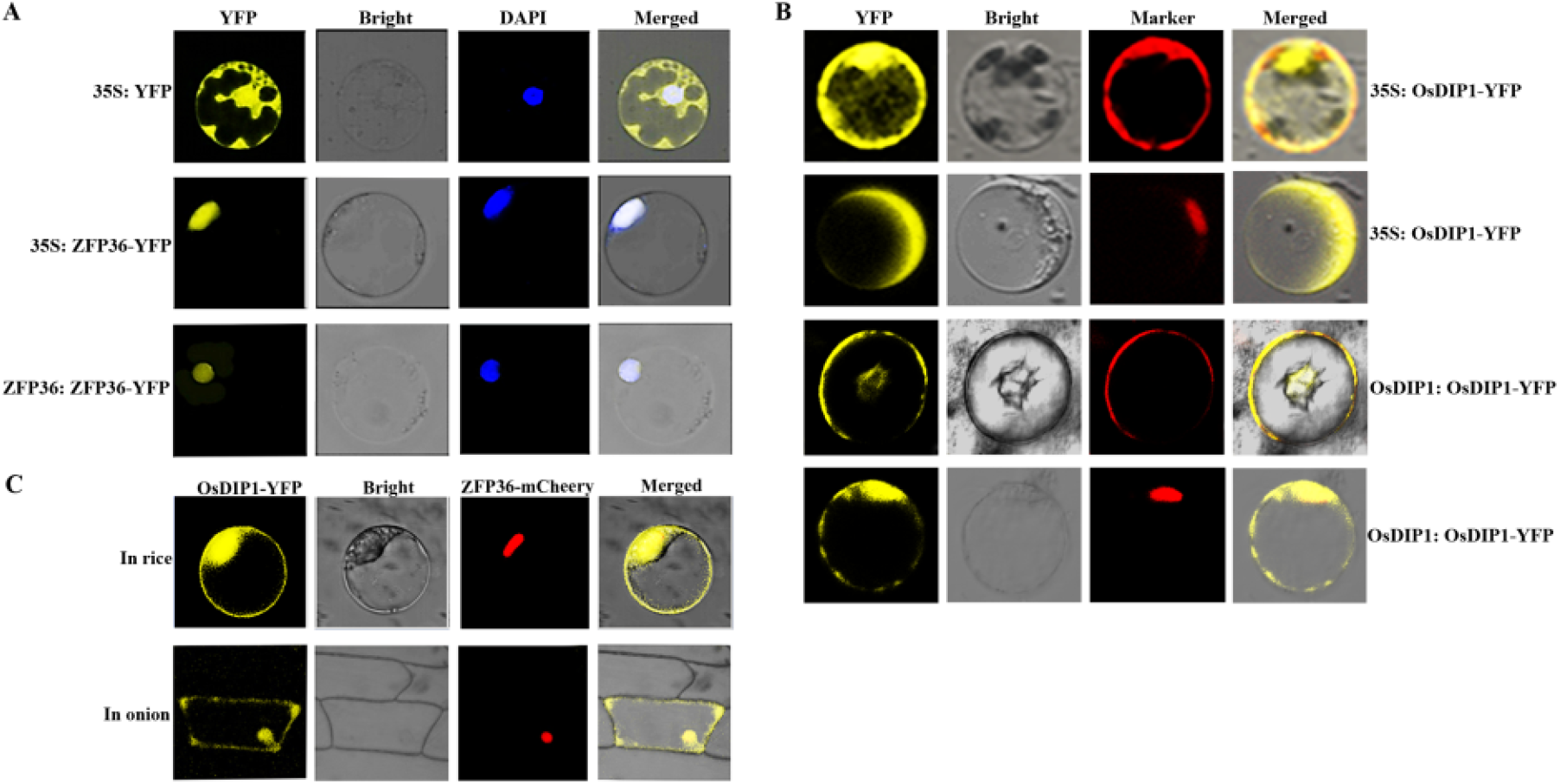
Subcellular localization of OsDIP1 and ZFP36, and its co-localization. (A) ZFP36 localization in rice protoplasts. 35: ZFP36-YFP (with 35S as a promoter) and ZFP36: ZFP36-YFP (with ZFP36 native promoter as a promoter) were separately transformed into rice protoplasts via PEG-mediated transient transformation, (4’,6-Diamidino-2-Phenylindole, Dihydrochloride) DAPI was used as the nuclear dye. (B) OsDIP1 localization in rice protoplasts. 35: OsDIP1-YFP (with 35S as a promoter) was co-expressed in the rice protoplasts with the membrane marker protein (row 1) and nuclear marker protein (row 2). OsDIP1: OsDIP1-YFP (with OsDIP1 native promoter as a promoter) was co-expressed in the rice protoplasts with the membrane marker protein (row 3) and nuclear marker protein (row 4). Membrane marker protein (MM), pm-rk CD1007 (Delporte et al. 2014); Nuclear marker protein (NM) red fluorescent protein (RFP)::H2A (Yu et al. 2015). (C) Co-localization of OsDIP1 and ZFP36 in onion. OsDIP1-YFP and ZFP36-mCheery were co-transformed into onion epidermal cells via GenGun Bombardment. Images were taken after incubation for 16 h.

**Fig. S2.**
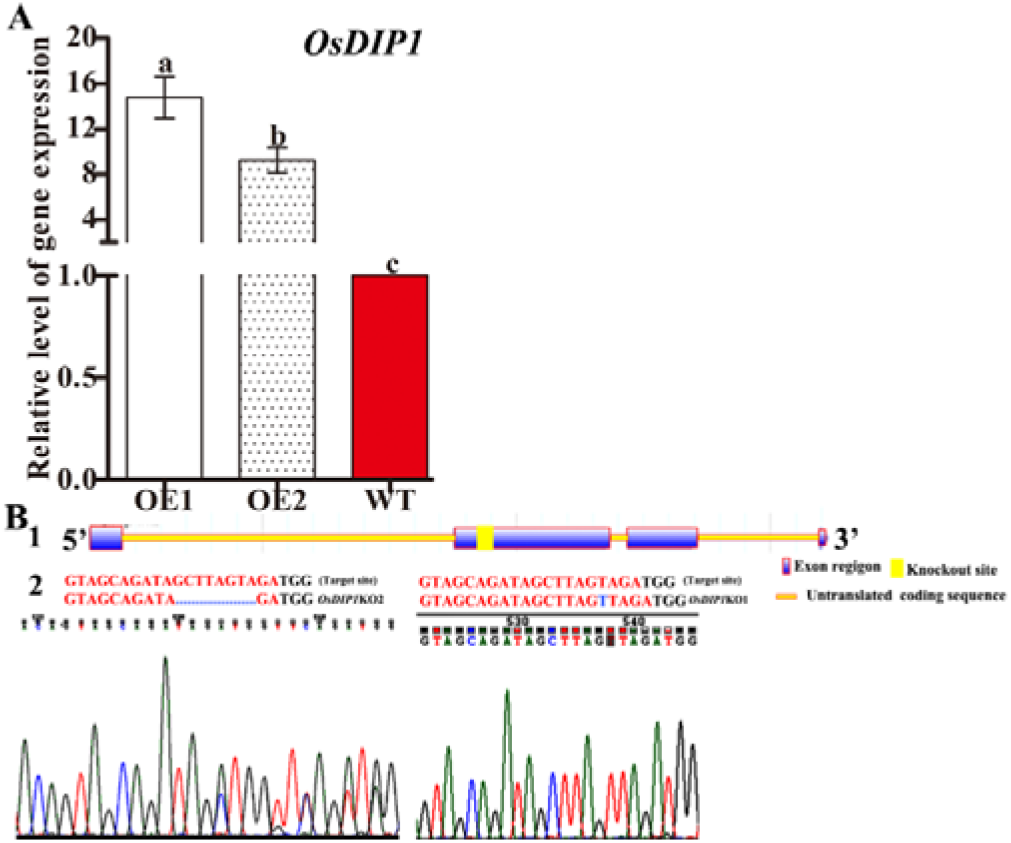
Expression level of *OsDIP1* mutants. (A) qRT-PCR analysis of the expression levels of *OsDIP1* in *OsDIP1*-overexpressed plants (OE1, OE2), and WT plants. (B) Identification the knockout site of Crispr/Cas9 mediated knockout of *OsDIP1* plants. (1) - the structure of OsDIP1 (1); (2) - the sequencing graph.

**Fig. S3.**
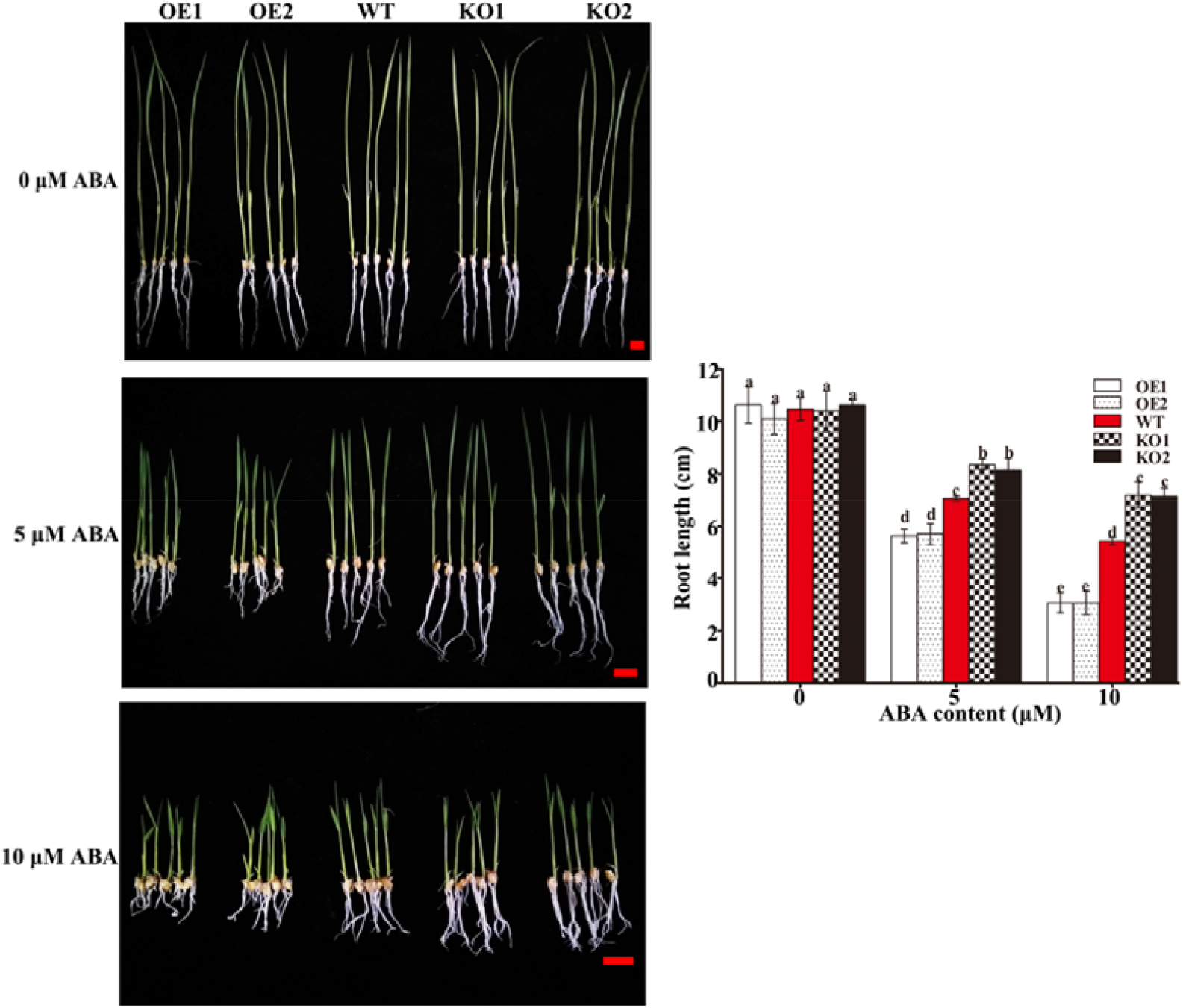
OsDIP1 participates in ABA response. The phenotype (left) and statistic analysis of root length (right) of rice plants under different content of ABA. 3-d-old *OsDIP1*-OE1, *OsDIP1*-OE2, *OsDIP1*-KO1, *OsDIP1*-KO2, and WT rice plants were treated with different concentration of ABA (0, 5, 10 μM) for 10 d. bar= 1 cm.

## Notes

### Competing Interest Statement

The authors have declared no competing interest.

